# Aspiration-assisted Freeform Bioprinting of Tissue Spheroids in a Yield-stress Gel

**DOI:** 10.1101/2020.05.31.122309

**Authors:** Bugra Ayan, Zhifeng Zhang, Nazmiye Celik, Kui Zhou, Yang Wu, Francesco Costanzo, Ibrahim T Ozbolat

## Abstract

Bioprinting of cellular aggregates, such as tissue spheroids or organoids, in complex three-dimensional (3D) arrangements has been a major obstacle for scaffold-free fabrication of tissues and organs. In this research, we unveiled a new approach to the bioprinting of tissue spheroids in a yield stress granular gel, which exhibited unprecedented capabilities in freeform positioning of spheroids in 3D. Due to its Herschel-Bulkley and self-healing properties as well as its biological inertness, the granular gel supported both the positioning and self-assembly of tissue spheroids. We studied the underlying physical mechanism of the approach to elucidate the interactions between the aspirated spheroids and the gel’s yield-stress during the transfer of spheroids from cell media to the gel. We demonstrate the application of the proposed approach in the realization of various freeform shapes and self-assembly of human mesenchymal stem cell spheroids for the construction of cartilage and bone tissues.

## Introduction

Three-dimensional (3D) bioprinting within granular gels or suspension baths exhibiting Herschel-Bulkley or Bingham plastic properties has recently become a powerful approach to create complex-shaped anatomically-accurate tissues and organs^1–9^. Carbopol microgels have been one of the popular granular gel medium due to its shear thinning and self-healing properties, in which the granular gel transforms from a stable solid state into a flowing fluid phase when exposed to an external stress that exceeds its yield stress^6,10–12^. As the nozzle moves inside the granular gel, the gel locally fluidizes when in contact with the nozzle but then rapidly solidifies after the nozzle has passed thus supporting the bioprinted tissue constructs^1,7^. In most cases of bioprinting in a granular gel, cells are bioprinted while encapsulated within a hydrogel formulation, resulting in limited cell densities.^13^ Hence, cellular aggregates, such as tissue spheroids, possess greater promise due to their favorable properties in building native-like tissues^14–16^.

Bioprinting of spheroids is an attractive approach, in which spheroids are used as building blocks for fabrication of tissues that mimic the native counterparts in terms of histology and physiology^14,16,17^. Several spheroid bioprinting techniques have been reported. The first technique is extrusion-based bioprinting^18^, in which spheroids are loaded in a syringe barrel and extruded in a delivery gel medium one by one. However, spheroids self-assemble readily in the syringe and are prone to break apart during the extrusion process. Concurrently, support structures need to be 3D printed to facilitate the aggregation of extruded spheroids. An important advance has been made by utilizing the Kenzan method^19^, where spheroids are skewered on a needle array. Since the position of each spheroid depends on the needle size, location and arrangement, freeform (i.e., complex-shaped) bioprinting of spheroids is quite challenging as the spheroid positioning of spheroids along the z-axis (direction parallel to the needles) is not independent in each layer. Drop-on-demand bioprinting has also been reported to deposit spheroids^20^. In this approach, spheroids are encapsulated within gel droplets. As such, drop-on-demand bioprinting has inherent limitations on the precision of the 3D bioprinting process. To overcome some of major the challenges of current techniques, we recently demonstrated an aspiration-assisted bioprinting (AAB) technique^21^ enabling precise bioprinting of spheroids into or onto sacrificial or functional gel substrates. However, freeform bioprinting of spheroids in 3D has been a long-standing problem due to the layer-by-layer-building nature of the existing techniques.

Here, for the first time, we demonstrate the freeform bioprinting of tissue spheroids by precisely positioning them in a self-healing biologically-inert granular gel in 3D allowing for the subsequent self-assembly of the bioprinted spheroids towards fabrication of tissues and organs. We used our previously demonstrated AAB technique to aspirate and pick spheroids and, taking advantage of the Herschel-Bulkey properties of the granular gel receiving the spheroids, we succeded in the direct transfer of spheroids from the cell media and their freeform positioning within the granular gel on-demand. In order to better understand the response of biologics to the bioprinting process, we studied the underlying mechanism explaining interactions between the spheroids and the granular gel during bioprinting. We then explored the potential of our Aspiration-assisted Freefrom Bioprinting (AAfB) technique in building complex-shaped configurations and demonstrated multiple applications, including cartilage and bone tissues throughout this study.

## Results

### 2.1 Working mechanism of AAfB

In this study, we further advanced our recently published Aspiration-assisted Bioprinting (AAB) technique^21^ to demonstrate the freeform bioprinting of spheroids within a granular gel. Specifically, aspiration forces were used to pick up spheroids from the spheroid reservoir (placed inside the cell media compartment) and transfer them into the granular gel (occupying inside the support gel compartment) one by one (**Figs. 1A1–1A7**). The spheroids were transferred from the cell media through a highly mobile transition interface into the self-healing granular hydrogel. In general, gels have small elasticity and high viscosity, and their mechanical response is usually described by a viscoelastic model^22^. Here, we present some elementary moment balance arguments leading to the estimate of the minimum aspiration pressure that is needed for a spheroid to be transferred from the media to the gel compartment. With reference to **Figs. 1B1–1B2**, whether the spheroid was moving through the interface or through the gel, we observed that the spheroid was acted upon by forces due to its interaction with its environment and with the nozzle. If the aspiration pressure fell below a critical value *P_b_*, the spheroid would separate from the nozzle, typically by pivoting against the trailing edge of the nozzle (trailing relative to the direction of motion). We label the pivot point by *T* in **Fig. 1B1**. We denote by *F_R_* the magnitude of the resultant force acting on the spheroid due to its interaction with the environment. Referring to **Fig. 1B1**, we observed that at the critical pivot condition the only forces contributing to the moment about the point *T* are the resultant of the applied aspiration pressure distribution and the force with magnitude *F_R_*. With this in mind, we can then estimate the critical aspiration pressure *P_b_* by considering the balance of moments about *T*. Clearly, to proceed to such an estimate we need to know both the values of *F_R_* and its direction as well as the state of motion of the spheroid. Since *F_R_* represents the resistance offered by the gel to the spheroid’s motion, we make the simplifying assumption that, when the spheroid is moving at a constant speed along a horizontal line, the resistance is also horizontal and with a line of action going through the spheroid’s center. Under these simplified conditions, the moment balance about *T*, ∑*M_T_* = 0, yields the following relation:

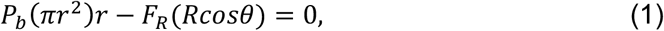

where, with reference to **Figs. 1B1–1B2**, *r* is the nozzle’s radius and *θ* is such that tan *θ = r/R*.

**Fig. 1:**
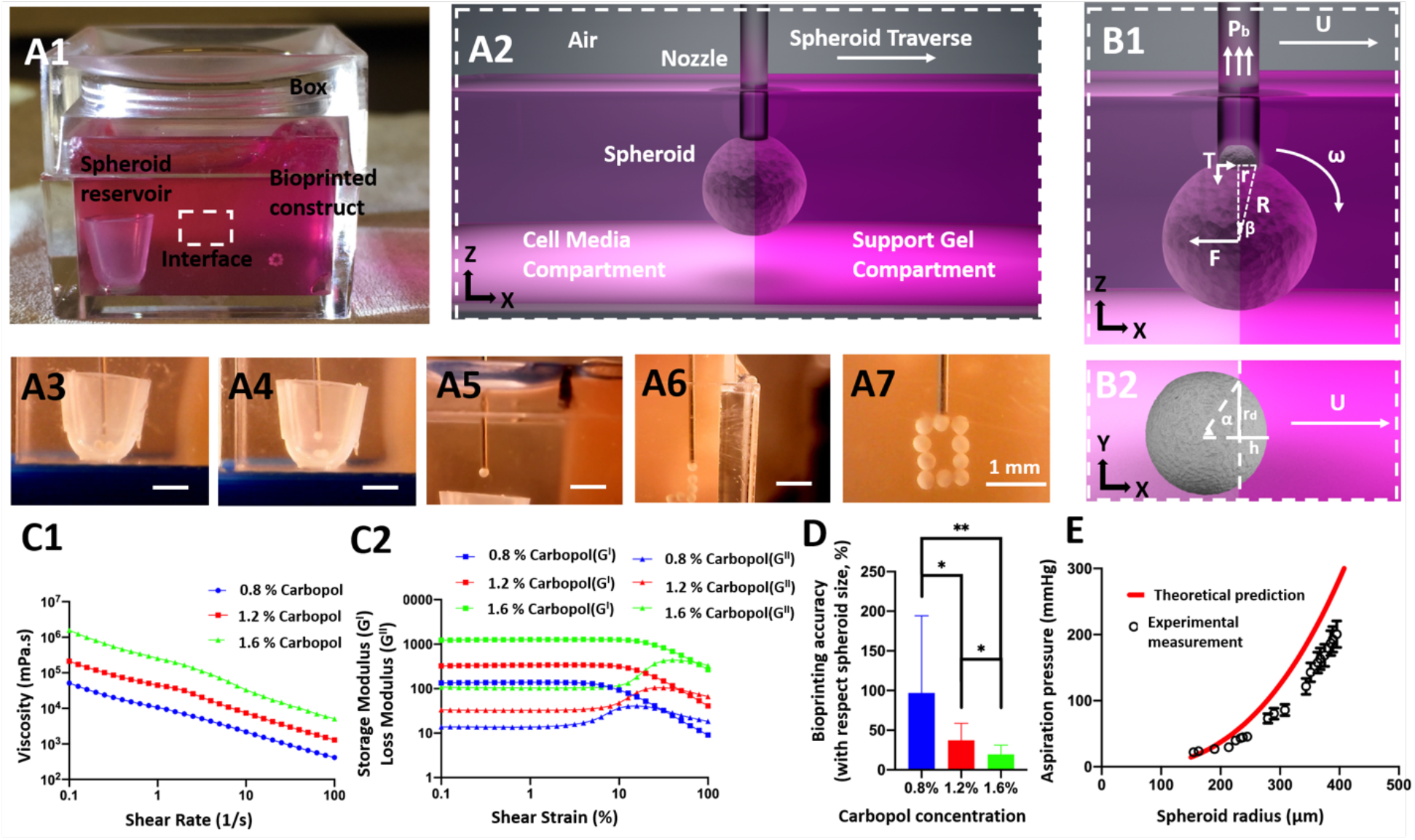
AAfB of spheroids in a yield stress gel. **(A1)** The bioprinting setup, where a box was filled with the yield-stress gel and cell media. **(A2)** A schematic showing spheroid traverse. **(A3-A7)** Real-time images showing a step-by-step illustration of the whole process, where spheroids were picked from the reservoir in the cell media and traversed the interface into support gel. **(B1-B2)** Schematics showing physical parameters involved in transferring of spheroids from the cell media to the support gel. **(C1-C2)** Rheological properties of the support gel (Carbopol) at different concentrations. **(D)** Bioprinting accuracy of the support gel at different concentrations (with respect to spheroid size) (*n*=5; \**P*<0.05 and \*\**P*<0.01). **(E)** Confirmation of the theoretical approach using the experimental validation for spheroids a ranging from 150 to 450 μm in radius bioprinted in 1.2% Carbopol gel. Note that human mesenchymal stem cells (MSCs) spheroids were utilized in all experiments. The theoretical relation is plotted according to Eq. (6).

Solving Eq. (1) for *P_b_*, we obtain

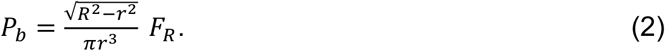

Next, we need to provide an estimate for the value of *F_R_*. This estimate can be complex in that *F_R_* is determined by different physics depending on the position of the spheroid relative to the interface between the medium and the gel compartments.

When the spheroid is moving through the gel as a constant speed, it is reasonable to assume that *F_R_* = *F_D_*, where *F_D_* is the drag acting on a sphere moving at a constant speed in a viscous fluid under laminar conditions. In fact, treating the spheroid as a rigid particle with a radius *R* (< 450 μm), the relevant Reynolds number^23^ is Re = 2*ρ_gel_*UR/*η*_0_, where *ρ_gel_* is the mass density of the gel, which is assumed to be the same as water (as a matter of fact, the mass density of the spheroids can also be assumed to be that of water: *ρ_s_* = *ρ_gel_* = *ρ_w_*), *U*~2.5 mm/s is the bioprinting speed (also the speed of the spheroid’s center-of-mass, and *η*_0_ is the gel’s Newtonian equivalent viscosity or zero-shear rate viscosity 44 Pa·s). Under these assumptions, *Re* is on the order of ~10^-6^ confirming that the flow around the spheroid during bioprinting is indeed laminar. Under these conditions, we can use the well-known formula *F_D_* = 6*πRUη_U_*, where the value of viscosity *η_L_* depends on *U* as the gel is shear-thinning ^24^.

More complex is the estimation of *F_R_* when the spheroid is traversing the interface between the medium compartment and the gel. In this case, we can distinguish four contributions to *F_R_*: again the drag exerted on the spheroid by its surroundings (*F_D_*), the resistance provided by the elasticity of the gel below the yield limit as the spheroid is indenting the gel (*F_E_*), the thermodynamic force (*F_I_*) representing capillary effect at the interface, and nonlinear and dynamic terms (*F_N-A_*), neglecting fluctuations and the rotational effects^25^, as the motion cannot be treated as being steady:

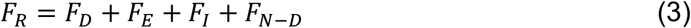

Whether in the gel or at the interphase, for simplicity, we will estimate *F_D_* using the same drag formula mentioned earlier scaled to account for the fact that the spheroid is not completely in the gel (**Fig. 1B2**): *F_D_ = Ur_d_η_U_*(6*α* + 8*sin α* + *sin* 2*α*), where *r_d_* is the contact radius and where the advancing angle *α* is defined via the relation tan *α* = *r_d_*/(*r_d_* — *h*), *h* being the indentation depth^24^ (**Fig. 1B2**). We feel that his estimate is acceptable in an effort to understand what physics dominates the value of *F_R_*. Clearly, the maximum resistance provided by the gel to the spheroid after traversing the interface is *FA = 6πr_d_Uη_u_*, as previously discussed. Referring to **Fig. 1C1,** our experimental rheological characterization indicates that the gel should be modeled as a (shearthinning) Herschel-Bulkey fluid with viscosity:

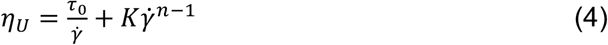

where *τ*_0_ is the yield stress, 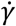 is the shear rate which we estimate as 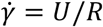 for the motion in the gel or as 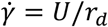 for the motion through the interface, *K* is the consistency index, and *n* is the power-law exponent (*n*<1 for shear-thinning fluids^26^). Two fluid property constant can be identified from the power-law shear-thinning regime in **Fig. 1C1**, *n* can be obtained by adding one to the slope of the viscosity versus shear rate curve and the consistency index *K* is equal to the viscosity of the gel when the shear rate is equal to 1. From our experiments, we see that *K*=44 (Pa·s^n^) and *n*=0.3. Thus, 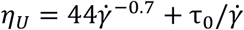, with a unit of Pa·s.

At the initial stage of contact^27^, *F_E_* = 4*πEhR* or using the same geometric configuration as above, *F_E_* = 4*πER*^2^(1 – cosα), *E* being the gel’s Young’s modulus. *E* can be estimated from rheological measurements of the storage shear modulus *G*’. Specifically, we have *E* ≈ 2*G*’(1 + *v*) = 3*G*’, where *v* is the Poisson ratio, which, for an incompressible material like Carbopol, can be taken to be equal to 0.5^28^. Our measurements of *G*’ are reported in **Fig. 1C2**. The term *F_E_* is only considered while the spheroid is traversing the interface and neglected when the spheroid is fully submerged in the gel.

Another term is the thermodynamic interfacial force is experienced when the spheroid is traversing the media-gel interface. The maximum value can be estimated to be:

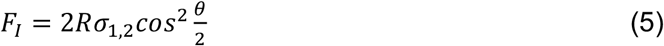

Here, *σ*_1,2_ is the surface tension coefficient between the media and gel. The last term, *F_N-A_*, includes the nonlinear and dynamic terms, such as rotation and inertia. Typical values for the surface tension coefficient are a few tens of mN/m. For the 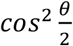 term, we assume an ideal value of 1 to maximize the interfacial effect. Considering the ratio of interfacial and drag term, 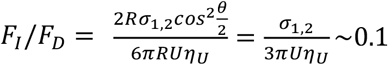.

Based on the above force analysis, the main contribution to the term *F_R_* is the drag experience by the spheroid as it moves through the gel, so that

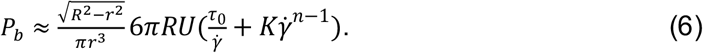

*η_U_* is the viscosity at printing speed of 2.5mm/s. As a result, *P_b_* is a function of *R, r, U*, gel properties (*K, n, τ*_0_). We have not included the terms that are a weak function of E, *σ*_1,2_ and *θ*, which are negligible compared to the viscosity of the gel. However, while crossing the interface, *F*_E_ term should also be included in the estimation of *P*_b_.

In order to determine an appropriate gel concentration for AAfB, we preferred to test 0.8, 1.2, and 1.6% concentrations of Carbopol, where such a range was comparable with respect to Carbopol concentration used in a previous study^13^. Our rheological experiments (**Fig. S1**) demonstrated a yield stress value of 5.3, 25.7, and 136.1 Pa for 0.8, 1.2, and 1.6% Carbopol, respectively (**Fig.1C2, Fig. S1**).

All concentrations showed shear-thinning properties indicated by decreasing viscosity with shear rate (**Fig. 1C1**), and solid to fluid transition occurred at ~13, 28, and 57% strain for 0.8%, 1.2% and 1.6% Carbopol, respectively. **Fig. 1D** shows bioprinting positional accuracy with respect to the spheroid size, which was improved with increasing Carbopol concentration such that 1.6, 1.2, and 0.8% Carbopol yielded 19, 37% and 97% positional accuracy, respectively. As shown by the error bars in **Fig. 1D**, the positional precision for 0.8, 1.2, and 1.6% concentrations were determined to be ~97, 22, and 12 %, respectively. In order to validate the theoretical approach, we performed bioprinting experiments to establish a relationship between *r* and *P_b_*. As indicated in **Fig. 1E,** the theoretical approach was confirmed by the experimental approach and the results were close to each other, particularly for spheroids with smaller radii. Yet, increasing aspiration pressure may lead to deformation of the spheroids^21^. Spheroids needed to be transferred in a safe manner without leading to significant deformations. As its know that external stressors induce considerable damage to cell viability^29^, 1.2% Carbopol concentration was preferred to be used in our further experiments.

### Applications of the AAfB technique

To demonstrate to the potential of our AAfB technique, we demonstrated the bioprinting of a DNA-strand **(Figs. 2A1–2A3)**, of the acronym PSU for Penn State University (**Figs. 2B1–2B2, and Supplementary Video 1**), and of five layers of circles forming a cylinder **(Figs. 2C1–2C4)** using uniform size human mesenchymal stem cell (MSC) spheroids (~175 μm in radius). We also bioprinted a double DNA-shaped strand MSC spheroids with different radii (150 and 450 μm). In addition to complexed-shaped configurations, we also demonstrated the AAfB of functional tissues, including cartilage and bone.

**Fig. 2:**
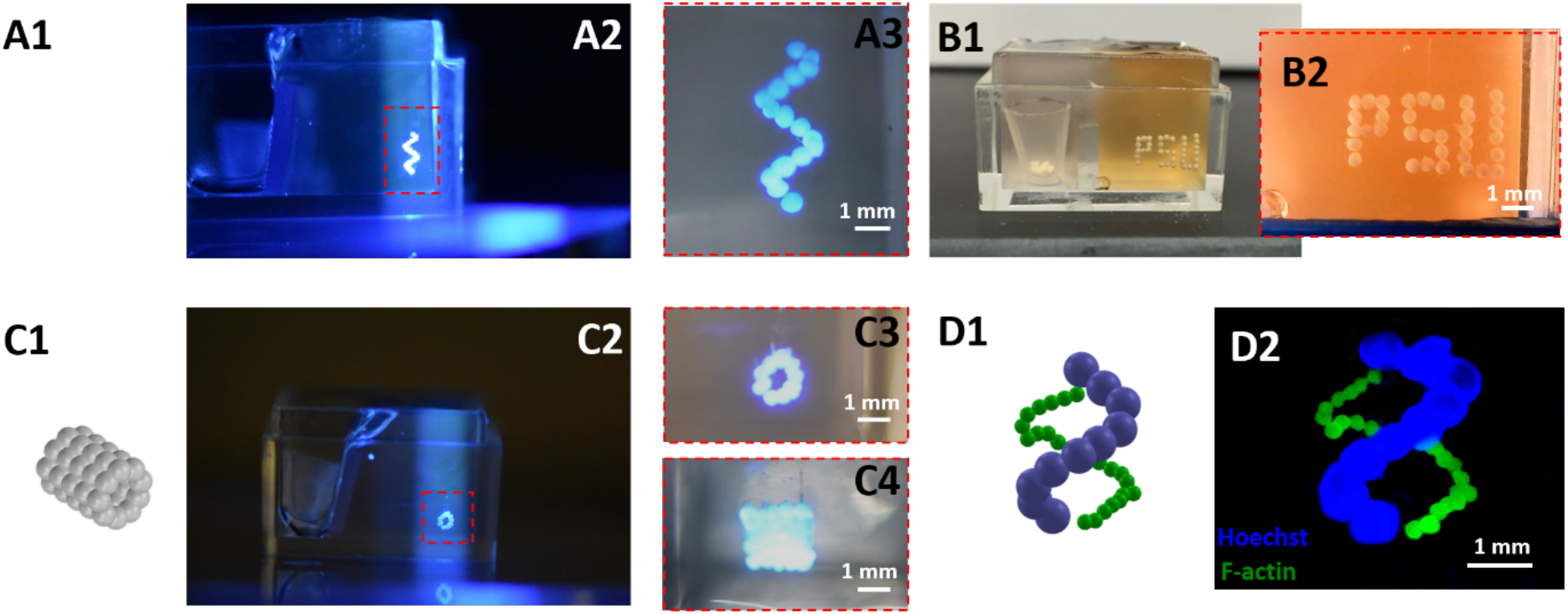
AAfB of spheroids in a Carbopol gel. Schematic illustration and photographs of 3D bioprinted **(A1-A3)** helix-shape (MSC spheroids), **(B1-B2)** initials of Penn State University (PSU, MSC spheroids), **(C1-C4)** 5-layer tubular (MSC spheroids), and **(D1-D2)** double helix-shape constructs using MSC spheroids with 150 μm (F-actin) and 450 μm (Hoechst) in radius.

Tubular cartilage tissues were bioprinted using MSC spheroids following two strategies in order to investigate the effect of the chondrogenic differentiation timeline on the functional and structural properties of bioprinted tissues **(Fig. S2)**. In the first strategy, which we will refer to as Strategy I, MSC spheroids were maintained in the growth media for three days and then were 3D bioprinted into a tube shape on Day 3. The bioprinted constructs were removed from Carbopol on Day 4 and further maintained in a chondrogenic induction medium for 20 days. In the second strategy, which we will refer to as Strategy II, MSC spheroids were maintained in the growth medium for three days followed by a 19 day culture in a chondrogenic induction medium, and finally bioprinted on Day 22. Upon sufficient fusion, the bioprinted constructs were removed from Carbopol gel on Day 23 and samples were collected for further analysis on Day 24. In order to understand the physical and biological properties of spheroids used in both strategies, we performed histological examinations (**Figs. 3A1–3A6**), size and surface tension measurement (**Figs. 3B1–3B2**), and protein quantification (sulfated glycosaminoglycan (sGAG) content) (**Fig. 3B3**).

**Fig. 3:**
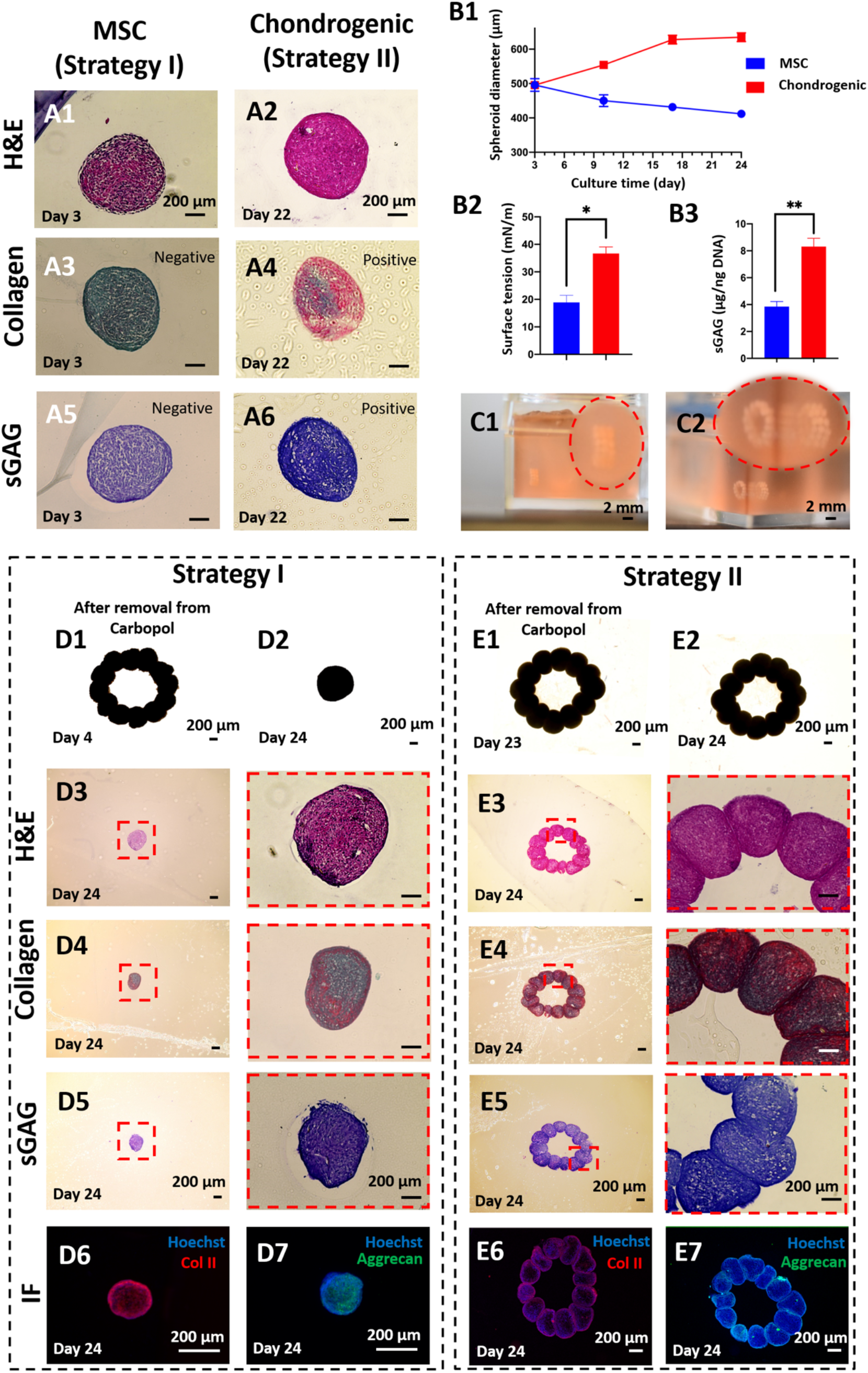
AAfB of tubular cartilage tissues. Histological staining of MSC (at Day 3, prior to bioprinting for Strategy I) and chondrogenic spheroids (at Day 22, prior to bioprinting for Strategy II) **(A1-A2)** H&E, **(A3-A4)** Picrosirius Red/Fast Green, and **(A5-A6)** Toluidine Blue staining. **(B1)** Radius change of MSC and chondrogenic spheroids over 24 days (*n*=10). Note that chondrogenic spheroids were cultured in MSC growth media for the first three days of culture. **(B2)** Surface tension and **(B3)** sGAG content measurements (normalized to DNA amount of MSC and chondrogenic spheroids at Day 24) (*n*=3, \**P*<0.05, and \*\**P*<0.01). **(C1-C2)** 3-layer bioprinted tubular cartilage constructs in Carbopol. Strategy I: tubular cartilage tissue were bioprinted using MSC spheroids (at Day 3). **(D1)** A microscopic image showing a bioprinted construct after removal from the support gel. **(D2)** The final shape of the bioprinted cartilage at Day 24. Histological and immunostaining images of the bioprinted tissues at Day 24 **(D3)** H&E, **(D4)** Picrosirius Red/Fast Green, **(D5)** Toluidine Blue, **(D6)** COL-II, and **(D7)** Aggrecan staining. Strategy II: tubular cartilage tissues were bioprinted using chondrogenic spheroids at Day 22. (**E1**) A microscopic image showing bioprinted construct after its removal from the support gel at Day 23. **(E2)** The final shape of the bioprinted cartilage tissue at Day 24. Histological and immunostaining images of the bioprinted at Day 24 **(E3)** H&E, **(E4)** Picrosirius Red/Fast Green, **(E5)** Toluidine Blue, **(E6)** COL-II, and **(E7)** Aggrecan.

We first investigated the differences in the spheroids used for Strategy I (3 day culture in a growth medium) and Strategy II (3 day culture in a growth medium followed by a 19 day culture in a chondrogenic induction medium) prior to bioprinting in terms of Hematoxylin and Eosin (H&E) staining as well as collagen and sGAG content. MSC spheroids used for Strategy I were less dense (from H&E staining) and were negative for collagen and sGAG, whereas the spheroids used for Strategy II (after a 22 day culture in a chondrogenic medium) were larger in size, more dense, and were positive for Collagen and sGAG. Thus, in the rest of the study, we will refer to the spheroids used in Strategy I and Strategy II as MSCs and chondrogenic, respectively. We also traced the change in spheroid size during the 24-day culture time (**Fig. 3B1**). The radius of chondrogenic spheroids increased from 250 μm (on Day 3) to a value slightly larger than 300 μm (on Day 18), and retained their size for the remaining period of the culture until Day 24. The radius of MSC spheroids gradually decreased from 250 μm (on Day 3) to 200 μm (on Day 24). The surface tension is an important parameter that determines the structural integrity of the spheroids. The higher the surface tension the better the bioprinting will be due to the spheroids’ decreased sensitivity to aspiration forces^21^. In this regard, chondrogenic spheroids had a surface tension that was approximately twice that of MSC spheroids (**Fig. 3B2**). Furthermore, the surface tension values for both spheroids were within feasible ranges for bioprinting^21^. Finally, we also observed a 2.2-fold increase in the sGAG content (μg/ng DNA) for chondrogenic spheroids as compared to MSC spheroids (**Fig. 3B3**). We then used these spheroids from Strategy I and Strategy II to bioprint tubular cartilage tissues, following the corresponding culture protocol for each strategy **(Figs. 3C1–3C2)**. The bioprinted tubular shape was preserved during 1-day culture in Carbopol post-bioprinting and after removal of the tissue from the Carbopol **(Fig. 3D1)**. However, the tubular organization collapsed into a dense ball after 20 days of culture in the chondrogenic induction medium for Strategy I whereas the tube shape was preserved in Strategy II. We performed H&E, collagen and sGAG staining as well as immunofluorescent (IF) staining for Collagen II (Col II) and Aggrecan on bioprinted tissues on Day 24. In Strategy I, H&E staining showed compact arrangement of the bioprinted tissues **(Fig. 3D3)**. This finding was different from that reveal by the morphology and histology of MSC spheroids **(Figs. 3A1–3A2)**. In this case, collagen deposition was concentrated around the outer surface of the bioprinted tissues, and sGAG staining was positive **(Figs. 3D4–3D5)**. IF staining results showed that both Col II and Aggrecan staining were positive **(Figs. 3D6–3D7)**. Here, we showed that the bioprinted tissues in Strategy I exhibited chondrogenic properties; however, the bioprinted shape could not be retained because of the compaction of MSC spheroids. In Strategy II, we observed that the spheroids retained their shape, showed sufficient fusion between them, and retained original tubular arrangements (**Figs. 3E1-E2**). H&E staining was similar to that of the chondrogenic spheroids (**Fig. 3E3**). Collagen and sGAG staining were both positive, similarly to the case with chondrogenic spheroid (**Figs. 3E4-E5**). Again, IF staining showed positive Col II and Aggrecan (**Figs. 3E6-E7**).

We also demonstrated the bioprinting of bone tissue using osteogenic spheroids as building blocks. Osteogenic spheroids were fabricated in three different groups from MSCs, and their differentiation was characterized in detail **(Fig. S3)**. In Group 1, MSC spheroids were formed on Day 0 and cultured in a osteogenic differentiation medium for 28 days. In Group 2, spheroids were formed after MSCs were cultured on the tissue culture plate (TCP) for seven days, followed by an additional 21 days in osteogenic differentiation media. In Group 3, MSCs were culture on TCP in osteogenic differentiation media for 12 days before spheroids were formed. Spheroids were then cultured in a osteogenic differentiation medium for additional 16 days. For each group, spheroids were collected for analysis purposes on Days 14 and 28.

When the spheroids were compared on Day 28, H&E staining for Group 3 showed considerable more bone matrix deposition as compared to Groups 1 and 2 (**Fig. 4A1-A3**). Confocal images of Group 3 demonstrated the strongest expression of OSTERIX, which is a late-stage osteogenic differentiation marker (**Fig. S4)**. Expression of osteogenic genes were investigated for different groups of spheroids, including bone morphogenic protein-4 (BMP-4), osteocalcin (OCN), Collagen I (COL-1), bone sialoprotein (BSP), and OSTERIX at Days 14 and 28. Overall, all genes for all groups showed greater amount of expression on Day 28 as compared to Day 14. Although gene expressions on Day 14 exhibited no significant difference among groups, expression of BMP-4 (4.8- and 32.3-fold), COL-1 (3.6- and 30.5-fold), BSP (3.5- and 22.8-fold) and OSTERIX (5.3- and 37-fold) in Group 3 on Day 28 was significantly higher than those for Groups 1 and 2, respectively.

**Fig. 4:**
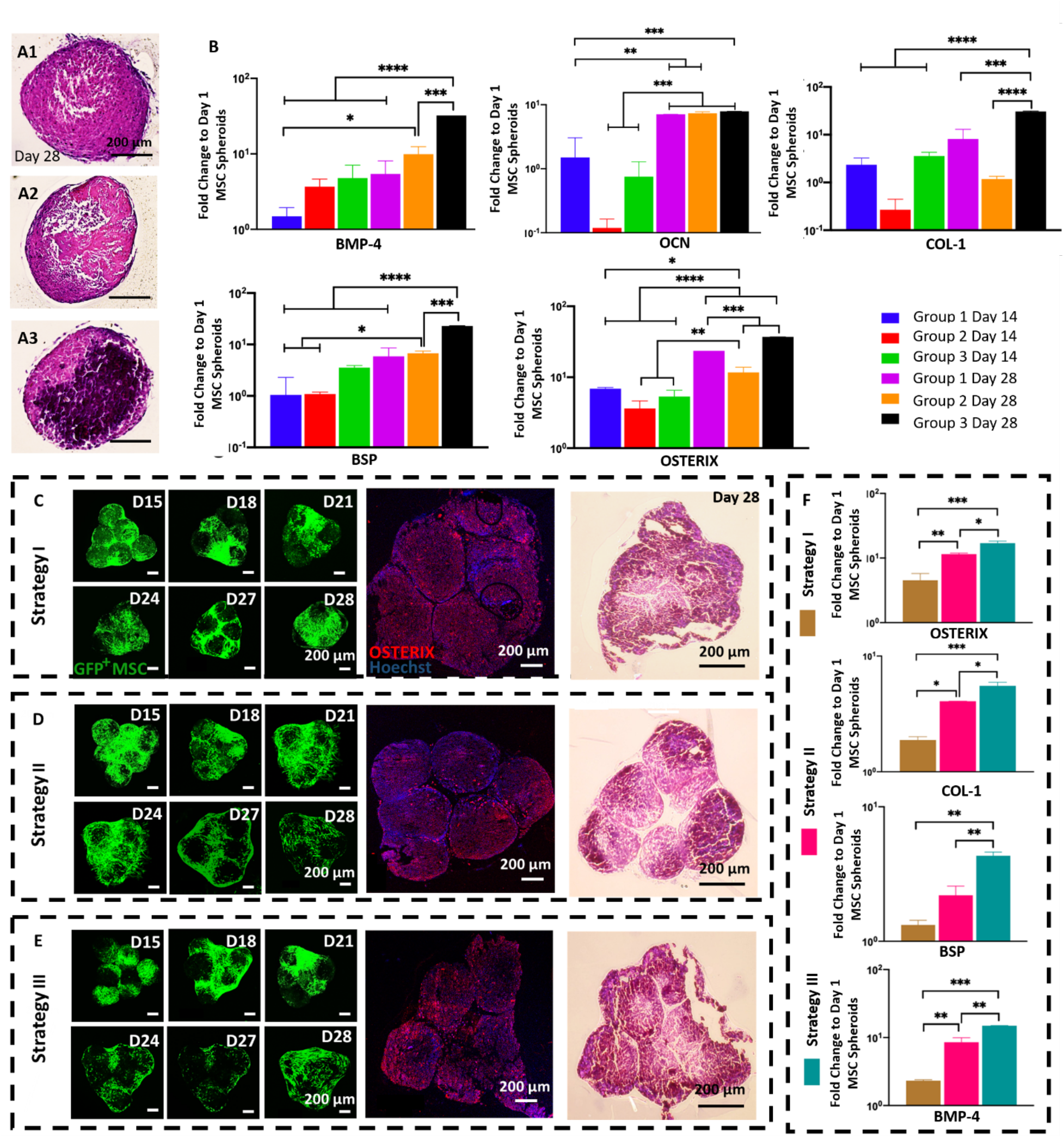
AAfB of osteogenic tissues. H&E images of spheroids of **(A1)** Group 1, **(A2)** Group 2, and **(A3)** Group 3 at Day 28. **(B)** *BMP-4, OCN, COL-1, BSP, and OSTERIX* gene expressions of Group 1 Day 14, Group2 Day 14, Group 3 Day 14, Group 1 Day 28, Group 2 Day 28,and Group 3 Day 28 *(n* = 3; \**P* < 0.05, \*\**P* < 0.01, \*\*\**P* < 0.001, and \*\*\*\**P* < 0.0001). Time-lapse images of GFP^+^ MSCs and their immune and H&E staining for bioprinted bone tissue using (C) Strategy I, (D) Strategy II, and (E) Strategy III. **(F)** *OSTERIX, COL1, BSP,* and *BMP-4* gene expressions of 3D bioprinted bone tissues cultured using different strategies *(n* = 3; \**P* < 0.05, \*\**P* < 0.01, and \*\*\**P* < 0.001).

For bioprinting of bone tissues, we followed three strategies in order to understand the role of the osteogenic induction timeline on the formation of bone tissue **(Fig. S5)**. In Strategy I, Group 1 day 14 osteogenic spheroids were used (i.e., MSC spheroids that were formed at Day 0 and then cultured in osteogenic differentiation media for 14 days). Group 1 spheroids were bioprinted on Day 14, and the tissue was removed from the Carbopol on Day 15, and cultured in osteogenic induction media for 13 days, completing a 28-day period in total. In Strategy II, Group 2 osteogenic spheroids were used (spheroids formed after 7-day 2D differentiation followed by 7-day 3D differentiation). Spheroids were bioprinted on Day 14 and the bioprinted tissues were removed from the Carbopol after spheroids fused each other sufficiently on Day 15, followed by 13 days of culture in a osteogenic induction medium. Finally, Strategy III utilized Group 3 osteogenic spheroids (spheroids formed after 12-day 2D differentiation followed by 2-day 3D differentiation). In this group, bioprinted tissues were cultured in the osteogenic differentiation media for 13 days after removal from the Carbopol.

We bioprinted triangle-shaped osteogenic tissues using six spheroids following these three Strategies. Fluorescent images of the tissues (at Days 15, 18, 21, 24, 27, and 28) were used to observe the shape changes due to the fusion and compaction of green fluorescent protein (GFP^+^) MSCs (**Figs. 4C–4E**). In Strategy I, the original shape could not be conserved due to compaction whereas in Strategies 2 and 3, the triangle-shape was well preserved. IF staining results showed that all of the three groups were positive for OSTERIX (**Figs. 4C–4E**). H&E staining also showed that the shape of the tissues in Strategy II and III were more triangular as compared to Strategy I. Expression osteogenic genes, including OSTERIX, COL-1, BSP, and BMP-4, were also evaluated. Constructs in Strategy III exhibited the highest expression level for all genes than those in Strategies I and II, namely 12.5- and 5.5-fold increase for OSTERIX, 3.7- and 1.5-fold increase for COL-1,3- and 2-fold increase for BSP and 12.6- and 6.4-fold increase for BMP-4, respectively. In addition, the expression level of OSTERIX (7-fold increase), COL-1 (2.2-fold increase) and BMP-4 (6.2-fold increase) were significantly higher in Strategy II as compared to those in Strategy I. Our results indicate that the longer the cells are exposed to induction media on 2D, the more pronounced the osteogenic differentiation in spheroids as well as bioprinted tissues.

## Discussion

Although extrusion-based bioprinting in granular gels has already been demonstrated in the literature^1,5–7,10,13,30^, its utilization in bioprinting of prefabricated cellular aggregates is quite challenging. Here, we presented a novel approach with the ability to bioprint cellular aggregates such as tissue spheroids in an accurate and precise manner in 3D. In this study, the presented AAfB approach enabled the freeform biofabrication of 3D complex-shaped constructs using spheroids as building blocks: we want to stress that this is not similarly achievable using existing bioprinting methods^15,16,19^. In addition to its strength in positioning of spheroids in 3D, the AAfB approach also made it possible to bioprint spheroids with a wide range of sizes (**Fig. 2**). While in this study we only utilized spheroids with radii ranging from 150 to 450 μm, we could readily modify the system to enable the bioprinting of spheroids with dimensions ranging from 100 μm up to almost 1 mm.

Bioprinting positional accuracy increased with the concentration of Carbopol. Due to the shear thinning behavior of the granular gel, when the Carbopol concentration was low (e.g., 0.8%), the granular gel liquefied and maintained insufficient viscosity and self-healing properties to hold the bioprinted spheroids in place accurately. The positional accuracy was increased with increasing Carbopol concentration. However, higher levels of aspiration pressure is required in these cases to transfer the spheroids from their initial location to their final placement. The higher aspiration pressure might induce substantial spheroid damage, such as their breakage during transition into the gel or their complete aspiration into the nozzle. Consequently, to exploit the potential of this new technique, it is crucial to determine of optimal gel properties and bioprinting speeds to guarantee the spheroids’ accurate placement while preserving their integrity and viability. Thus, Carbopol concentration of 1.2% was preferred to use throughout our experiments.

While it might be convenient to assume that the gel properties are uniform within the entire gel domain, our empirical observation is that the cell medium diffuses into the gel and changes the gel properties accordingly. In particular, we note that the gel properties changed considerably when the bioprinting time was prolonged. This said, we managed to bioprint larger constructs by minimizing the total volume of medium in order to control the issues related to the diffusion induced gel heterogeneity. Clearly, the bioprinting time could be minimized by increasing the bioprinting speed. However, a substantial increase in the bioprinting speed could also result in failure as spheroids could easily get stuck at the medium/gel interface due to the substantial resistance exerted by the gel (**Supplementary Video 2**). In this regard, we used 2.5 mm/s as our preferred bioprinting speed, which allowed for a rapid enough assembly of the presented tissue models while remaining safe enough to successfully transfer the spheroids from the cell media to the gel. Meanwhile, we also maintained an amount of medium sufficient to support the growth and viability of the bioprinted tissues. Also, we kept the bioprinted tissues in the Carbopol gel for only one day as one day was sufficient to induce partial fusion of spheroids and provide necessary structural integrity. Although we attempted to minimize the issues that could be encountered due to the biological inertness of the Carbopol as well as the possible risk of pH changes^13,31,32^ (which could bring severe damages to the viability of bioprinted constructs if not controlled otherwise), we experienced the disassembly of the bioprinted cartilage and bone tissues quite a few times during the removal of those tissues from Carbopol. This could be due to the change in the pH level during *in vitro* incubation impairing the biological and physical properties of Carbopol resulting in limited fusion between spheroids. In general, controlling the pH level of Carbopol and removing the fused spheroids from the Carbopol was not trivial^1,11,13^. Thus, synthesis and development of novel support gels, possessing optimal mechanical properties in terms of yield stress, shear thinning as well as additional physical properties such as self-healing, transparency, biocompatibility, and the ability to be drained from the bioprinted tissue, will greatly improve the deployment of platforms for fabrication of scalable human tissues and organs at the clinically-relevant volumes. The theoretical estimation of the force exerted on the spheroid form its environment during bioprinting was elementary and meant to capture how the nozzle aspiration pressure scales with the printing speed and the spheroid’s radius. Our model is particularly limited in capturing the details of the spheroid transfer through the medium/gel interface where the elastic and plastic behavior of the gel both significantly contribute to the force on the spheroid. Finally, our modeling did not include any considerations on the deformability of the spheroids, which is of primary concern when assessing viability. The future enhancement of the proposed bioprinting technique, especially when trying to tune the gel’s physical properties to achieve an optimal printing accuracy and viability, can benefit from a more sophisticated analysis of spheroid motion mechanics.

In our attempts preceding this study, we aspirated and lifted spheroids using a glass pipette with a radius of about 40 μm (see our recently published work^21^). We encountered problems in the use of a pipette when we transitioned spheroids in the gel domain, i.e., as we crossed the medium/gel interface. As depicted in **Supplementary Video 3**, spheroids were prone to bounce at the pipette tip because of insufficient aspiration forces against the drag force, which was due to the reduced exposure area of aspiration. In addition, as the pipette enlarged significantly toward its upper portion, we observed other issues such as substantial damages to the gel along with slower and diminished healing. Because of these reasons, we switched to metallic straight nozzles with a larger nozzle radius (inner radius of 100 μm). As long as we bioprinted spheroids with a radius of at least 150 μm at least, the metallic straight nozzles proved sufficient to perform the presented bioprinting work. This said, smaller nozzle tips or even pipette tips could still be utilized for bioprinting of spheroids with radii smaller than 50 μm.

In this study, chondrogenic spheroids were bioprinted into a tubular arrangement using two different strategies in order to understand the role of MSC or chondrogenic spheroids in successful bioprinting of tubular cartilage tissues. We identified considerable differences between MSC and chondrogenic spheroids in term of biological, structural, and mechanical properties. In particular, MSC spheroids shrank in size while chondrogenic spheroids grew over time, which could be due to the significant deposition of chondrogenesis-related extracellular-matrix (ECM) deposition, which in turn yielded higher surface tension and sGAG content in chondrogenic spheroids. Bioprinting of chondrogenicly differentiated spheroids generated tissues with improved chondrogenic properties and shape fidelity.

In bioprinting of osteogenic spheroids, three different strategies were designed to study the role of monolayer versus 3D induction on successful formation of bone tissue with controlled morphology. The results indicated that longer culture period in monolayer improved the construct fidelity as evidence by the result of Strategy III (**Fig. 4E**). This could be due to the increase exposure of MSCs to osteogenic differentiation media (REF) or improved osteogenesis of MSCs due to the substrate stiffness of TCP^33,34^. As MSCs in 3D spheroid culture had limited integrin-mediated adhesion with respect to TCPs, we observed enhanced bone formation at the gene and protein level^33^. It is also known that, osteogenicly differentiated MSCs could have limited proliferation, which might reduce the fusion and compaction of osteogenic spheroids in bioprinted bone tissues. Along with successful bioprinted outcomes, we also encountered some failures which we believe can be overcome with improved gel properties and pH control, or the use of other yield-stress gels.

In sum, we presented a highly effective approach in 3D bioprinting and positioning of tissue spheroids by explaining the interplay between the bioprinting process and gel yield-stress. Such a platform enabled us to pattern tissue spheroids with a high degree of geometric complexity, which will have tremendous applications such as, but not limited to, tissue engineering and regenerative medicine, disease modeling, drug screening, and biophysics.

## 3. Materials and method

### 3.1 Preparation of the granular hydrogel

To prepare the granular gel, 0.8, 1.2 and 1.6% (w/v) Carbopol ETD 2020 NF (Lubrizol Corporation, OH) were dispersed in human chondrogenic or osteogenic differentiation media (Cell Applications, CA) under sterile conditions. NaOH was added drop-wise to the Carbopol-dispersed gel to adjust the pH to 7.4, which facilitated the maximum swelling of Carbopol and its biocompatibility. The GH was then homogenized using a vortex blender for 20 min, centrifuged at 1000 x g for 15 min, and incubated at 37°C and 5% CO_2_ before further use.

### 3.2 Fabrication and differentiation of MSC spheroids

Human mesenchymal stem cells (MSCs, RoosterBio Inc., Frederick, MD) and green fluorescent protein (GFP)-labeled MSC (Cyagen, CA) were used in experiments. Both types of MSCs were cultured in RoosterBasal™ MSC medium, composed of RoosterBooster™ MSC-XF growth supplement, 100 U/mL penicillin, 100 μg/mL streptomycin, and 2.5 μg/mL fungizone (Life Technologies, CA), under a humidified atmosphere with 5% CO2 at 37 °C. To prepare spheroids, MSCs were trypsinized and centrifuged to form cell pellets. 200 μL of the cell suspension (2.5×10^5^-2×10^6^ cells/mL) were transferred into each well of a 96-well plate (Greiner Bio One, NC). Cells were cultured in MSC growth media during spheroid formation, and the medium was changed every three days. MSC spheroids were differentiated into chondrogenic and osteogenic lineages for different applications using human chondrocyte and osteoblast differentiation media, respectively (Cell Applications, CA).

### 3.3 Rheological analysis of the granular gel

Rheological measurements of the granular gel were performed using a MCR 302 rheometer (Anton Paar, VA) using a 25 mm diameter parallel-plate geometry. A Peltier system was employed for temperature control. Amplitude tests were applied to determine viscous and elastic properties of the gel at a constant frequency of 1 Hz and a strain range from 0.01 to 100% at a constant temperature of 25 °C.

### 3.4 AAfB process

For the bioprinting setup, we utilized our previously developed AAB system ^21^, a square petri dish was used to hold the gel and cell culture media (**Fig. 1A**), where a Polydimethylsiloxane (PDMS) slice was used to transfer and confine the gel in order to obtain a vertically-oriented interface. Spheroids were placed in a reservoir which was submerged in the tissue-specific media. When the reservoir was transferred into the Petri dish, tissue specific cell media was filled to cover the remaining area in the Petri dish. A 27G needle (Nordson, OH) was used to pick the spheroids from the reservoir and to transfer them from cell culture media into GH, with a speed of 2.5 mm/s. Two microscopic cameras (one for each side of the Petri dish) were used to visualize the bioprinting process in real time. In order to validate the theoretical results, MSC spheroids with a wide range of radius (from 150 to 400 micron) were bioprinted into 1.2% Carbopol medium (GH).

### 3.5 Accuracy and precision measurement of bioprinting

In order to investigate the effect of Carbopol concentration on the positional accuracy and precision, MSC spheroids were printed at pre-determined target positions in 0.8, 1.2 and 1.6 % Carbopol according to the protocol described in our previous work^21^. A calibration slide (Motic, China) was placed at the bottom of the petri dish to monitor the target position. After spheroid deposition, images of the bioprinted spheroids (*n*=5) were taken using cameras under the calibration slide and analyzed using ImageJ software (National Institutes of Health, MD). Accuracy was represented as the root mean square error (RMSE), which was calculated using the equation as below:

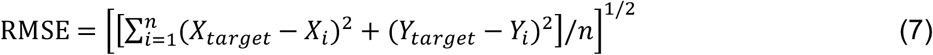

where *X_target_* and *Y_target_* represent X and Y coordinates of the target position, respectively, *X_i_* and *Y*-are the position of the measured values in X- and Y-axis, respectively, and *n* is the sample size. Precision was represented as the square root of the standard deviation.

### 3.6 Physical properties of MSC and chondrogenic spheroids

The chondrogenic differentiation of MSC spheroids was started on Day 3, and the radius of spheroids were measured by an EVOS^®^ microscope (Invitrogen, MA) until Day 24. Surface tension of spheroids was also measured according to the protocol described in our previous published work^35^. Briefly, customized straight micropipettes (~40 μm in radius), fabricated from glass pipettes (VWR, PA) using a P2000 Flaming/Brown micropipette puller (Sutter Instrument, CA), were used to aspirate spheroids, which were monitored via a STC-MC33USB monochromatic camera (Sentech, Japan). Surface tension of the MSC and chondrogenic spheroids were measured on Day 24.

### 3.7 Histological analysis of spheroids

MSC and chondrogenic spheroids were fixed with 4% paraformaldehyde and sectioned with paraffin embedding to obtain 10 μm sections. Hematoxylin and eosin (H&E) was performed on the sections using Leica Autostainer XL (Leica, Germany). Sections were also stained with Picrosirius Red to detect collagen content. The reagent was prepared by dissolving 0.1% Direct Red 80 and 0.1% Fast Green FCF in saturated aqueous picric acid (1.2% picric acid in water, Sigma Aldrich, MO). The solution was then applied to sections and incubated for 20 min. For sGAG visualization using Toluidine Blue O staining, sections were incubated in a Toluidine Blue solution (0.1% in DI water, Sigma Aldrich, MO) at room temperature for 2 min. The dye was then removed, and samples were washed twice with DI water, followed by dehydration with ascending alcohol and clearing with xylene. All samples were mounted and imaged using the EVOS^®^ microscope.

### 3.8 Evaluation of sGAG content

sGAG content was determined by DMMB dye-binding assay. MSC and chondrogenic spheroids were washed and digested in 500 μL solution of 0.1 mg/mL papain extraction reagent at 65 °C in water bath for 18 h. 20 μL of the digested samples were mixed with 200 μL DMMB solution and the absorbance was measured at 525 nm using a microplate reader (PowerWaveX, BioTek, Winooski, VT). Serially diluted solution of chondroitin 4 sulfate was prepared as the standard and the sGAG content was calculated according to the standard curve. The DNA content of same samples was also measured using the Quant-iT™ PicoGreen dsDNA Assay Kit (Molecular Probes Inc., Eugene, OR) according to the manufacturer’s instructions. Fluorescence intensity was determined by a SpectraMax multidetection microplate reader (Molecular Devices, Inc., Sunnyvale, CA) using a wavelength of 480 nm (excitation) and 520 nm (emission). sGAG content from each sample was normalized to dsDNA content.

### 3.9 Bioprinting of tubular cartilage tissue

In order to optimize bioprinted tubular cartilage tissue, two strategies were designed. In Strategy I, MSC spheroids on Day 3 were used for bioprinting, and the bioprinted constructs were incubated for chondrogenic differentiation for another 21 days. In Strategy II, MSC spheroids were cultured with chondrogenic media for 19 days, followed by bioprinting and cultured for 2 days in the form of constructs. After bioprinting of chondrogenic spheroids, the excess amount of Carbopol was gently removed (without affecting the structural integrity of the bioprinted constructs) in order to maximize the diffusion of cell media to better support the growth of the tissue. Constructs obtained by two strategies were characterized by H&E staining, Picrosirius Red staining and Toluidine Blue O staining as described in Section 3.7 to visualize the tissue morphology and chondrogenesis.

### 3.10 Immunohistochemistry of the bioprinted cartilage tissues

Primary monoclonal antibodies were purchased from Abcam (MA) and fluorescence-conjugated secondary antibodies were purchased from Life Technologies (CA). Sections of MSC and chondrogenic spheroids were treated using Triton-X 100 (0.1 % in PBS) for 10 min and blocked with normal goat serum (NGS, 10 % in PBS) for 1 h. Samples were then incubated with monoclonal rabbit anti-human collagen type II (COL-II, 1:200), mouse anti-human aggrecan (1:50) and NGS (negative control) for 1 h, respectively. Samples were washed twice with PBS and incubated using secondary antibodies (goat anti-rabbit IgG (H+L)-Alexa Fluor 647 for COL-II, and goat anti-mouse IgG (H+L)-Alexa Fluor 488 for aggrecan, 1:200) for 1 h. Samples were also incubated with Hoechst 33258 (1:200) for 5 min. Images for each marker were taken using a Zeiss Axiozoom microscope (Carl Zeiss Microscopy, LLC, Germany).

### 3.11 Gene expression of osteogenic spheroids using quantitative real-time polymerase chain reaction (RT-qPCR)

In order to investigate the effect of different strategies on the osteogenesis of spheroids, 3 groups were designed. In Strategy I, spheroids were prepared from MSCs and cultured for 28 days in osteogenic differentiation media. In Strategy ii, MSCs were cultured in monolayer for seven days, followed by fabricating and culturing spheroids for 21 days in osteogenic differentiation media. In Strategy III, MSCs were cultured in monolayer for 12 days, followed by fabricating and culturing spheroids for 16 days in osteogenic induction media. For all groups, the total induction period in monolayer culture and in the form of spheroids was kept 28 days in total.

For testing of bone-specific gene expression using RT-qPCR, single differentiated spheroids per sample were homogenized in TRIzol reagent (Life Technologies, CA), followed by adding 0.2 mL chloroform per 1 mL TRIzol reagent and centrifuging the mixture at 12,000 x g for 15 min at 4°C. The upper aqueous phase with RNA was transferred and RNA was then precipitated by adding 0.5 mL isopropyl alcohol per 1 mL TRIzol reagent, followed by centrifuging at 12,000 x g for 10 min, at 4°C. Subsequently, the precipitated RNA was rinsed twice by 75% ethanol, air-dried for 10 min and dissolved in 50 μL diethyl pyrocarbonate (DEPC)-treated water. RNA concentration was measured using a Nanodrop (Thermo Fisher Scientific, PA). Reverse transcription was performed using AccuPower^®^ CycleScript RT PreMix (BIONEER, Korea) following the manufacturer’s instructions. Gene expression was analyzed quantitatively with SYBR Green (Thermo Fisher Scientific, PA) using a QuantStudio 3 PCR system (Thermo Fisher Scientific). Bone-specific genes tested included OSTERIX (Transcription factor Sp7), COL-1, OCN (osteocalcin), BMP-4 (Bone Morphogenetic protein-4) and BSP (Bone sialoprotein). The reader is refereed to Table 1 for the gene sequences. Expression levels for each gene were then normalized to glyceraldehyde 3-phosphate dehydrogenase (GAPDH). The fold change of hMSCs spheroids after formation on Day 2 was set as 1-fold and values in osteogenic groups were normalized with respect to that of the group.

**Table 1.**
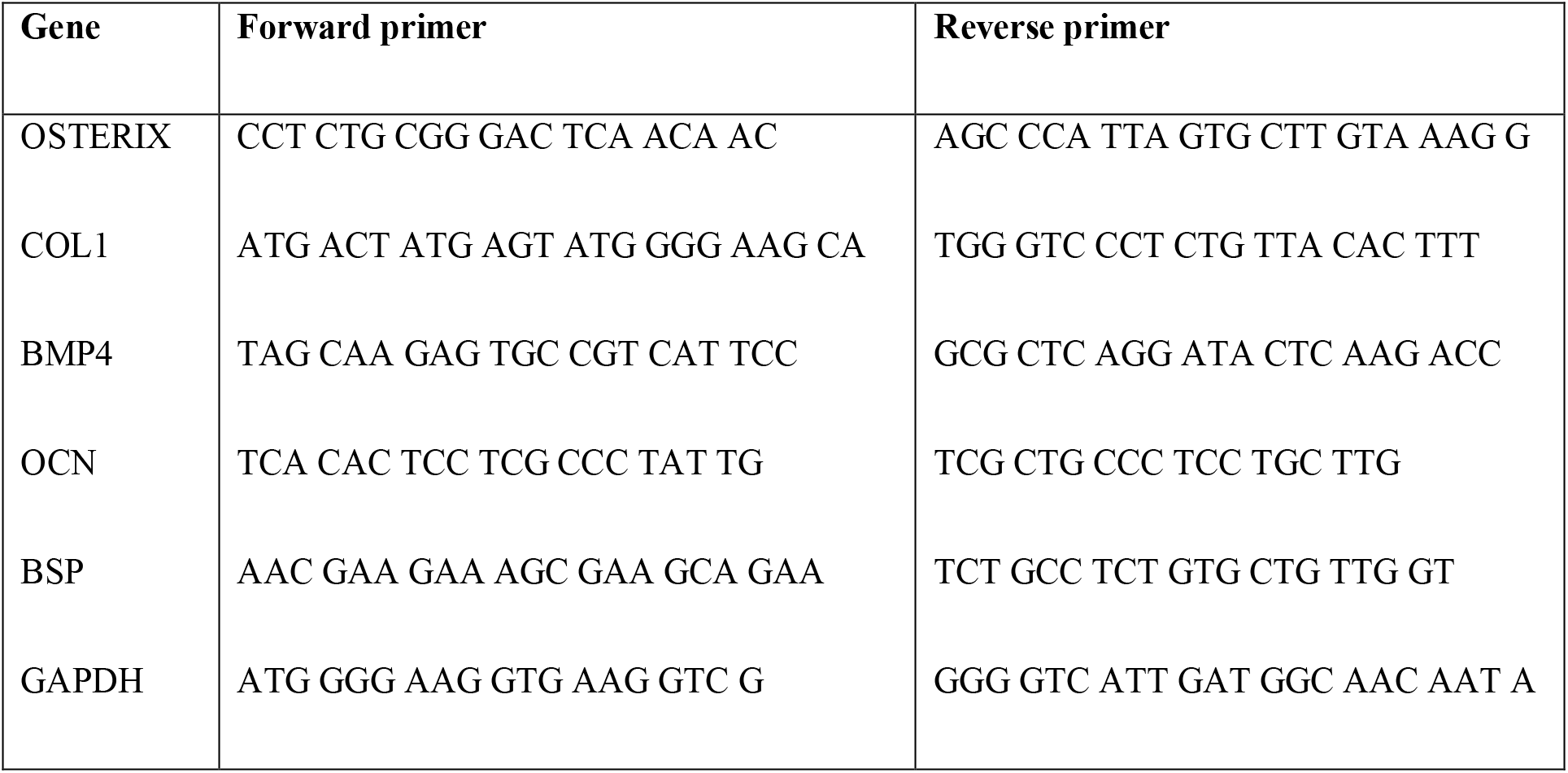
Primers of the measured mRNA for RT-qPCR

### 3.12 Bioprinting of osteogenic tissues

In order to investigate the effect of different osteogenic strategies on formation of bioprinted bone tissue, 3 groups were designed. In Group 1, spheroids were prepared using MSCs and cultured with osteogenic induction media for 14 days. In Group 2, MSCs were cultured with osteogenic induction in monolayer for seven days, followed by fabricating and culturing spheroids for seven days with osteogenic differentiation media. In Group 3, MSCs were cultured with osteogenic induction in monolayer for 12 days, followed by fabricating and culturing spheroids for two days with osteogenic induction media. After bioprinting of osteogenic spheroids, the excess amount of Carbopol was gently removed (without affecting the structural integrity of the bioprinted constructs) in order to maximize the diffusion of cell media to better support the growth of the tissue. For all groups, the total differentiation period in monolayer culture and in the form of spheroids were kept 14 days in total. Triangle bone structure were then bioprinted and cultured for 14 days in osteogenic differentiation media for a total of 28 days culture for all groups.

### 3.13 Biological characterization of bioprinted bone tissues

Immunohistochemistry staining was performed as explained in Section 3.10. To visualize latestage osteogenic formation, anti-Sp7/OSTERIX primary antibody (1:500 in 2.5% NGS) and goat anti-rabbit Alexa Fluor 568 secondary antibody (1:200 in 2.5% NGS) were used. Samples were imaged using a Zeiss LSM 880 Airyscan Confocal microscope (Zeiss, Oberkochen, Germany). RT-qPCR of bioprinted tissues were conducted as described in Section 3.11. H&E staining was carried out to visualize the morphology as described in Section 3.7.

### 3.14 Statistical analysis

All values were presented as mean ± standard deviation. Multiple comparisons were analyzed by using one-way analysis of variance (ANOVA) by Post-hoc Tukey’s multiple-comparison test was used to determine the individual differences among the groups. Differences were considered significant at *P < 0.05, **P < 0.01, and ***P < 0.001, and ****P < 0.0001. All statistical analysis was performed by Statistical Product and Service Solutions software (SPSS, IBM, USA).

## Acknowledgements

This work has been supported by National Science Foundation Award #1914885, Osteology Foundation Award #15-042, and a contract from The Lysosomal and Rare Disorders Research and Treatment Center Inc. We thank RoosterBio for providing MSCs and growth media. Authors thank D.N. Branford from PSU for assistance in designing the graphics in Fig.1. N.C. acknowledges the support from the Turkish Ministry of National Education.

## Contributions

B.A. and I.T.O. designed the research. B.A., Z.Z., F.C., and I.T.O. completed the theoretical work. B.A., N.C., K.Z., and Y.W. performed the experiments. All authors contributed to writing of the manuscript and agreed the final content of the manuscript.

## References

1. Bhattacharjee, T. et al. Writing in the granular gel medium. Sci. Adv. 1, e1500655–e1500655 (2015).

2. Mattsson, J. et al. Soft colloids make strong glasses. Nature 462, 83–86 (2009).

3. Saunders, B. R. & Vincent, B. Microgel particles as model colloids: theory, properties and applications. Adv. Colloid Interface Sci. 80, 1–25 (1999).

4. Dimitriou, C. J., Ewoldt, R. H. & McKinley, G. H. Describing and prescribing the constitutive response of yield stress fluids using large amplitude oscillatory shear stress (LAOStress). J. Rheol. (N. Y. N. Y). 57, 27–70 (2013).

5. Jeon, O. et al. Individual cell-only bioink and photocurable supporting medium for 3D printing and generation of engineered tissues with complex geometries. Mater. Horizons 6, 1625–1631 (2019).

6. Hinton, T. J., Hudson, A., Pusch, K., Lee, A. & Feinberg, A. W. 3D Printing PDMS Elastomer in a Hydrophilic Support Bath via Freeform Reversible Embedding. ACS Biomater. Sci. Eng. 2, 1781–1786 (2016).

7. Hinton, T. J. et al. Three-dimensional printing of complex biological structures by freeform reversible embedding of suspended hydrogels. Sci. Adv. 1, (2015).

8. Highley, C. B., Song, K. H., Daly, A. C. & Burdick, J. A. Jammed Microgel Inks for 3D Printing Applications. Adv. Sci. 6, 1801076 (2019).

9. McCormack, A., Highley, C. B., Leslie, N. R. & Melchels, F. P. W. 3D Printing in Suspension Baths: Keeping the Promises of Bioprinting Afloat. Trends Biotechnol. 38, 584–593 (2020).

10. O’Bryan, C. S. et al. Self-assembled micro-organogels for 3D printing silicone structures. Sci. Adv. 3, e1602800 (2017).

11. Ozbolat, V., Dey, M., Ayan, B. & Ozbolat, I. T. Extrusion-based printing of sacrificial Carbopol ink for fabrication of microfluidic devices. Biofabrication 11, 034101 (2019).

12. Jin, Y., Compaan, A., Bhattacharjee, T. & Huang, Y. Granular gel support-enabled extrusion of three-dimensional alginate and cellular structures. Biofabrication 8, 025016 (2016).

13. Bhattacharjee, T. et al. Liquid-like Solids Support Cells in 3D. ACS Biomater. Sci. Eng. 2, 1787–1795 (2016).

14. Mironov, V., Boland, T., Trusk, T., Forgacs, G. & Markwald, R. R. Organ printing: Computer-aided jet-based 3D tissue engineering. Trends in Biotechnology 21, 157–161 (2003).

15. Norotte, C., Marga, F. S., Niklason, L. E. & Forgacs, G. Scaffold-free vascular tissue engineering using bioprinting. Biomaterials 30, 5910–5917 (2009).

16. Mironov, V. et al. Organ printing: Tissue spheroids as building blocks. Biomaterials 30, 2164–2174 (2009).

17. Datta, P., Ayan, B. & Ozbolat, I. T. Bioprinting for vascular and vascularized tissue biofabrication. Acta Biomater. 51, 1–20 (2017).

18. Jakab, K. et al. Tissue Engineering by Self-Assembly of Cells Printed into Topologically Defined Structures. Tissue Eng. Part A 14, 413–421 (2008).

19. Moldovan, N. I., Hibino, N. & Nakayama, K. Principles of the Kenzan Method for Robotic Cell Spheroid-Based Three-Dimensional Bioprinting. Tissue Eng. Part B Rev. 23, 237–244 (2017).

20. Gutzweiler, L. et al. Large scale production and controlled deposition of single HUVEC spheroids for bioprinting applications. Biofabrication 9, 025027 (2017).

21. Ayan, B. et al. Aspiration-assisted bioprinting for precise positioning of biologics. Sci. Adv. 6, eaaw5111 (2020).

22. Style, R. W., Hyland, C., Boltyanskiy, R., Wettlaufer, J. S. & Dufresne, E. R. Surface tension and contact with soft elastic solids. Nat. Commun. 4, 2728 (2013).

23. Becker, L. E., McKinley, G. H., Rasmussen, H. K. & Hassager, O. The unsteady motion of a sphere in a viscoelastic fluid. J. Rheol. (N. Y. N. Y). 38, 377–403 (1994).

24. Sussman, M. & Smereka, P. Axisymmetric free boundary problems. J. Fluid Mech. 341, 269–294 (1997).

25. McKinley, G. H. Steady and transient motion of spherical particles in viscoelastic liquids. Transp. Process. Bubbles, Drops Part. 338–375 (2001).

26. Gabbanelli, S., Drazer, G. & Koplik, J. Lattice Boltzmann method for non-Newtonian (power-law) fluids. Phys. Rev. E 72, 046312 (2005).

27. Chakrabarti, A. & Chaudhury, M. K. Direct Measurement of the Surface Tension of a Soft Elastic Hydrogel: Exploration of Elastocapillary Instability in Adhesion. Langmuir 29, 6926–6935 (2013).

28. Tanner, R. I. Engineering Rheology. CEA, Chemical Engineering in Australia (Oxford University Press, 2000).

29. Yu, Y., Zhang, Y., Martin, J. A. & Ozbolat, I. T. Evaluation of cell viability and functionality in vessel-like bioprintable cell-laden tubular channels. J. Biomech. Eng. 135, 91011 (2013).

30. Heo, D. N. et al. 3D Bioprinting of Carbohydrazide-Modified Gelatin into Microparticle-Suspended Oxidized Alginate for the Fabrication of Complex-Shaped Tissue Constructs. ACS Appl. Mater. Interfaces (2020). doi:10.1021/acsami.0c05096

31. O’Bryan, C. S., Bhattacharjee, T., Marshall, S. L., Gregory Sawyer, W. & Angelini, T. E. Commercially available microgels for 3D bioprinting. Bioprinting 11, e00037 (2018).

32. O’Bryan, C. S., Kabb, C. P., Sumerlin, B. S. & Angelini, T. E. Jammed Polyelectrolyte Microgels for 3D Cell Culture Applications: Rheological Behavior with Added Salts. ACS Appl. Bio Mater. (2019). doi:10.1021/acsabm.8b00784

33. Shih, Y.-R. V, Tseng, K.-F., Lai, H.-Y., Lin, C.-H. & Lee, O. K. Matrix stiffness regulation of integrin-mediated mechanotransduction during osteogenic differentiation of human mesenchymal stem cells. J. Bone Miner. Res. 26, 730–738 (2011).

34. Duan, B., Yin, Z., Hockaday Kang, L., Magin, R. L. & Butcher, J. T. Active tissue stiffness modulation controls valve interstitial cell phenotype and osteogenic potential in 3D culture. Acta Biomater. 36, 42–54 (2016).

35. Hospodiuk, M. et al. Sprouting angiogenesis in engineered pseudo islets. Biofabrication 10, 035003 (2018).

