## Supplementary Information for "Aspiration-assisted Freeform Bioprinting of Tissue Spheroids in a Yield-stress Gel"

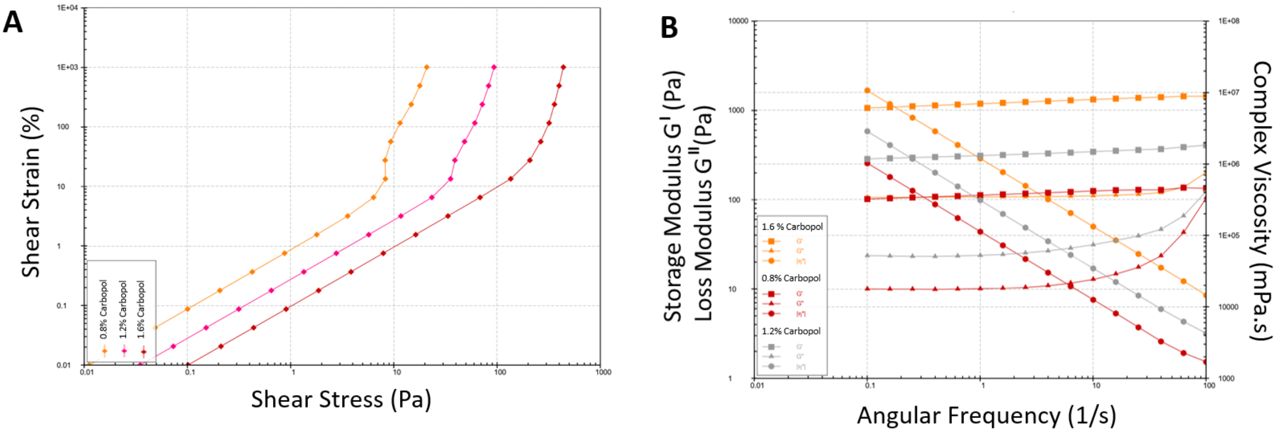


**Fig. S1: Rheological properties of Carbopol gel at different concentrations. (A)** Shear Strain- Shear stress **(B)** Frequency sweep of Carbopol gel at different concentrations.

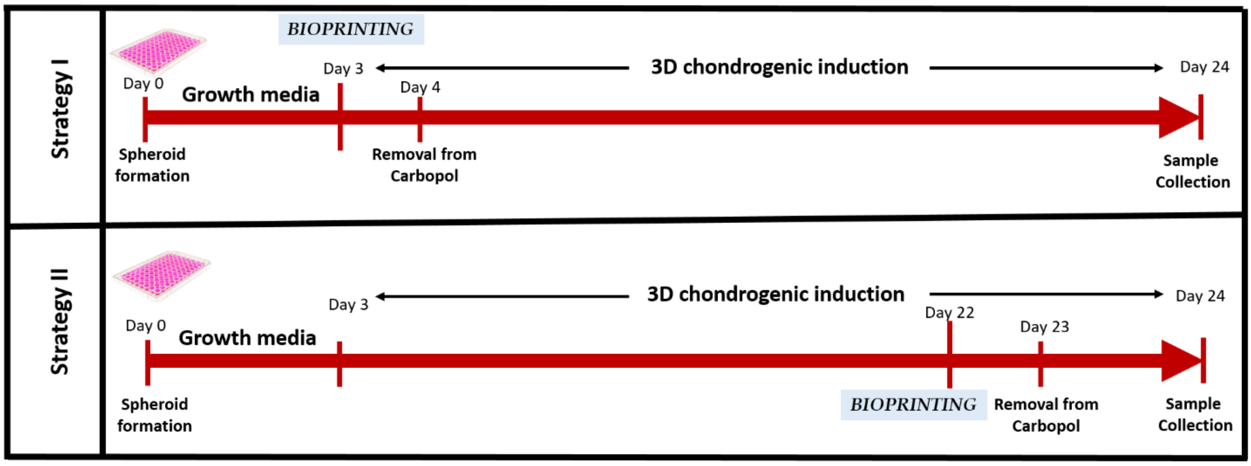


**Fig. S2: Culture strategies for tubular cartilage tissues, including Strategy I and Strategy II**

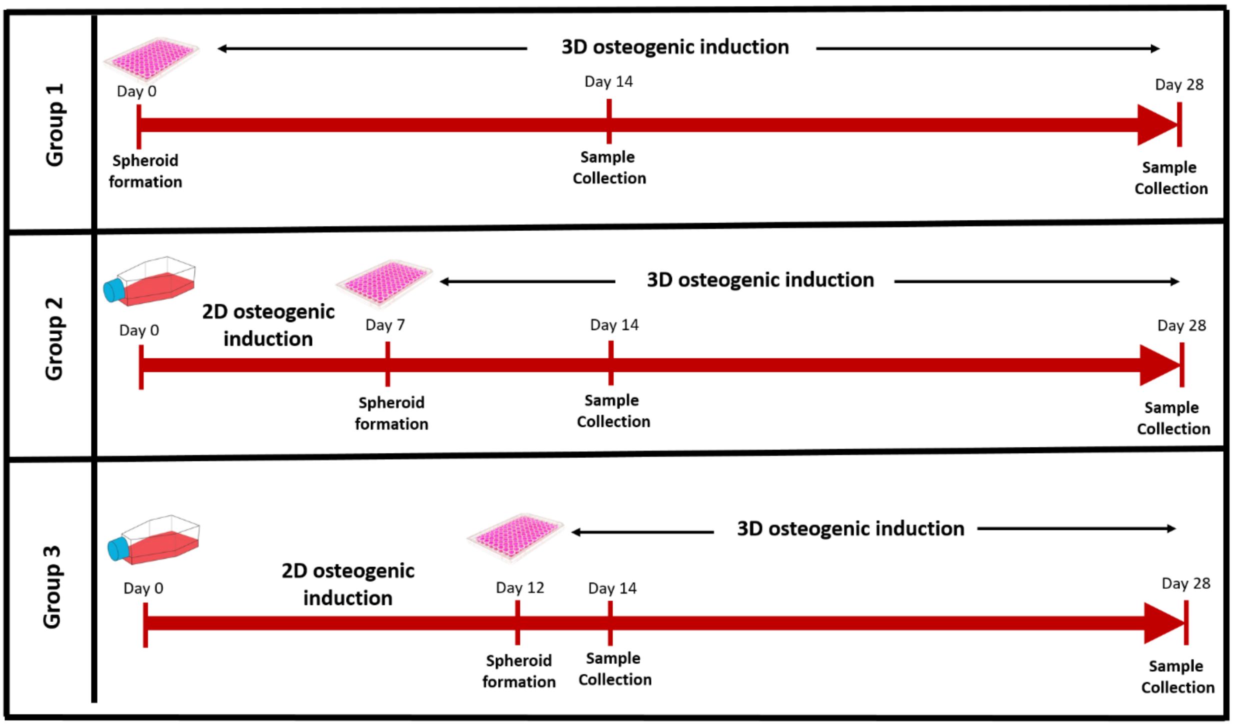


**Fig. S3: Timeline showing preparation and sample collection of osteogenicly-inducted MSC spheroids for Group 1, Group 2, and Group 3.**

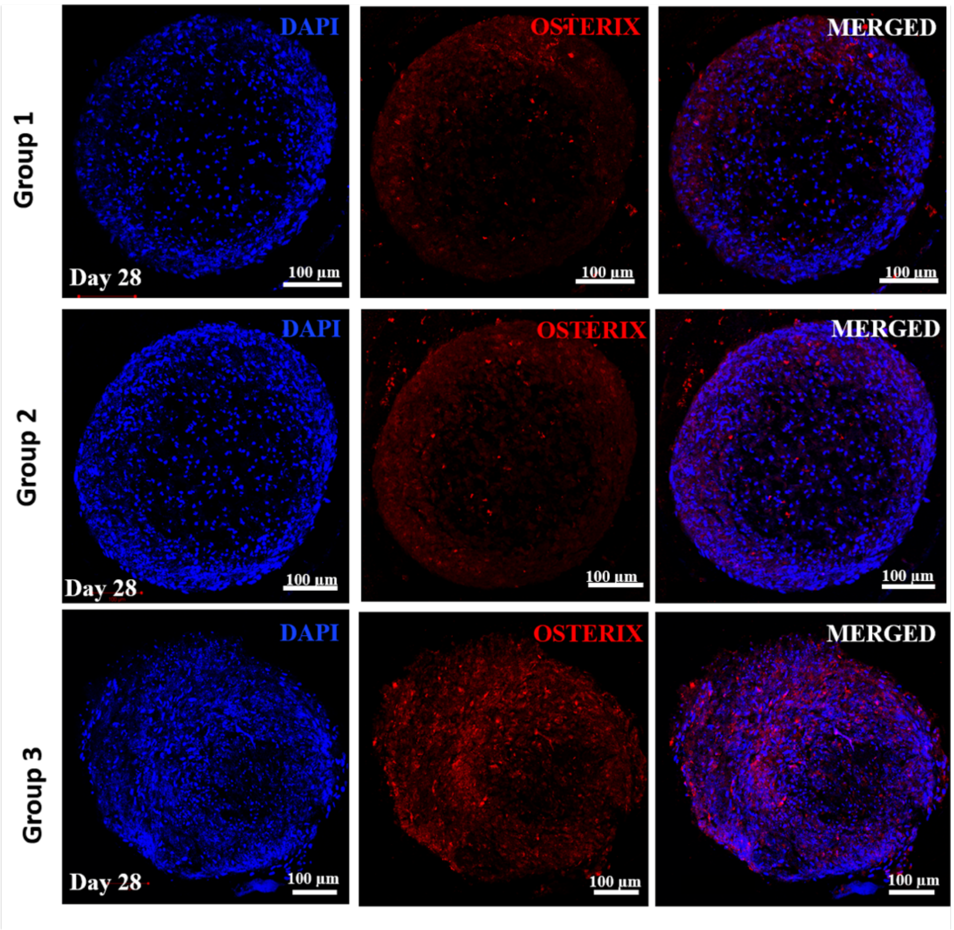


**Fig. S4: Confocal images of histological sections of different groups of osteogenic spheroids for DAPI, OSTERIX, and DAPI+OSTERIX at Day 28.**

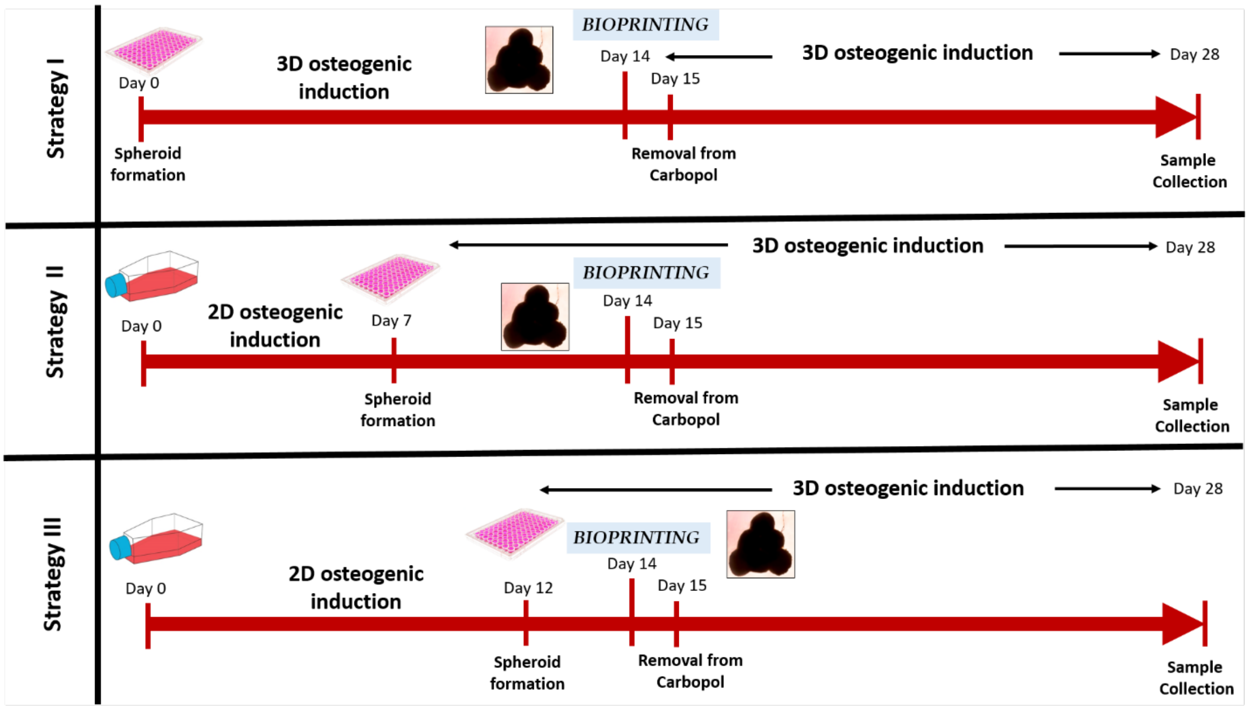


**Fig. S5: Culture strategies for bioprinted bone tissues including Strategy I, Strategy II, and Strategy III.**

**Supplementary Video 1: Bioprinting of spheroids in yield-stress Carbopol gel (2X Speed)**

**Supplementary Video 2: A spheroid got stacked at the interface during bioprinting process**

**Supplementary Video 3: Bouncing of a spheroid at the pipette tip while transitioning in Carbopol**
